# Suppression of optokinesis in the reafferent direction during pursuit eye movements

**DOI:** 10.1101/2024.09.16.613183

**Authors:** Omar Bachtoula, Melanie Ellul Miraval, Ignacio Serrano-Pedraza, David Souto

## Abstract

When tracking with the eyes an object moving against a textured background, the background retinal image moves in the opposite direction to the smooth pursuit eye movement. Optokinetic responses, such as optokinetic nystagmus (OKN) or ocular tracking, to this reafferent signal must be suppressed to sustain pursuit of the object-of-interest. We varied the contrast of a brief background motion to tell apart two plausible accounts of the suppression of optokinesis during pursuit; a visuomotor gain modulation account, which predicts that ocular tracking of background motion is suppressed in the same proportion at irrespective of contrast, and a sensory attenuation account, which predicts that larger contrasts are needed to elicit the same response. Unexpectedly, neither account fits ocular tracking in the reafferent signal direction. The combination of contrast-dependent gating, with maximal suppression observed with higher contrasts, and visuomotor gain modulation, provides a good fit for most observers’ data. Contrast-dependent gating promotes visuomotor stability in response to most salient signals, as a likely adaptation to the statistics of the environment.

**Significance statement:** For humans to be able to track small moving objects, there is a need for a mechanism to cancel optokinesis, that is reflexive eye movements towards prevalent visual motion. We show that this cancellation is not a simple “volume-control” reduction of responses to motion signals, as expected. This suppression also involves contrast-dependent gating, meaning that most salient signals are not allowed to modify the ongoing movement. This additional component could have arisen from an adaptation to image statistics of motion signals prevalent in natural environments.

## Introduction

In sensory systems, self-generated sensation, also known as reafference, must be accounted for to accurately signal external events^1,2^. A corollary discharge, a copy of the motor command conveying information about the impending movement to sensory brain areas^3^, could allow the recovery of external signals by discounting expected sensory consequences of movements. Corollary discharges are believed to play a role in maintaining perceptual continuity during saccades^4^, which can be achieved by different forms of downregulation of self-induced sensation^5,6^ (but see^7^). During saccades, self-generated retinal motion and changes in location are strongly attenuated by visual masking and by other active mechanisms^5^, contributing to the impression of visual stability despite discontinuous sensory input. In contrast, during smooth pursuit eye movements (pursuit for short), the attenuation of reafferent signals is less obvious and possibly unnecessary.

Pursuit eye movements, which aim to match a moving object’s velocity (e.g. a car), generate self-induced retinal motion from a textured background (e.g. a traffic scene). During pursuit, self-induced sensation may be used to calibrate motion perception^8^ or carry information that helps generate stable space-centric representations^9^. However, motoric responses to background retinal motion need nonetheless suppressing, since this reafferent signal is an ideal stimulus for optokinetic eye movements, such as OKN or ocular tracking, that conflicts with the ongoing voluntary pursuit eye movement.

Suppression of optokinesis during pursuit may take several forms, including the attenuation of self-generated sensation^6,10^. Several paradigms have been used to investigate the suppression of optokinesis during pursuit; such as by eliciting adaptation aftereffects or by eliciting reflexive response to sudden changes in the movement of the background^11,12^. The latter paradigm is well suited to test the contribution of sensory and pre-motor stages involved in the suppression of optokinesis.

Many neural responses, such as the firing rates of visual neurons^13^, or the amplitude of ocular tracking towards moving gratings^14^, increase with visual contrast or signal strength (e.g. proportion of coherent motion signals^10^) in a way that can be fit by the Naka-Rushton model^10^. This functional model, as illustrated in Figure 1, specifies the level at which responses asymptote, the rate at which they increase, and at which contrasts they increase. The model makes also clear predictions regarding the effects of sensory suppression and visuomotor suppression of optokinesis. If suppression occurs at the level of the response (response gain model), such as if a top-down signal from the frontal-eye-fields (FEF) inhibits pre-motor activity associated with responses to reafferent motion, we should have a proportional suppression in the amplitude at every signal level and, as a result, an increased maximal asymptotic response, as shown in Figure 1A. Response gain modulation would be consistent with the visuomotor gain modulation proposed^15,16^ to account for enhanced ocular responses to target trajectory perturbations during pursuit relative to fixation, where responses to visual motion are amplified to allow accurate pursuit in the presence of smaller retinal motion signals^17^. Response gain modulation could also be responsible for suppression of optokinesis observed in several studies and paradigms^11,12,18–20^.

**Figure 1.**
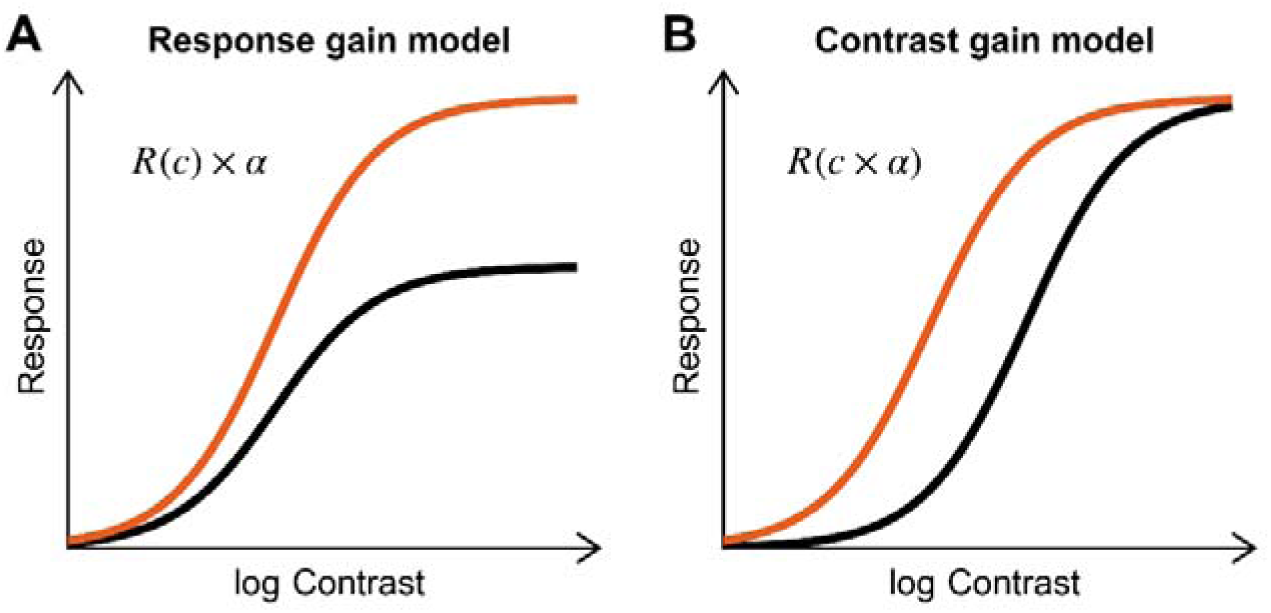
Illustrations of two fundamentally different ways in which a response could be suppressed. A) Response gain model refers to the amplification of a response. Amplifying the output of some neural process is equivalent to changing the asymptote (in the Naka-Rushton model) of the contrast-response function. B) The contrast gain model amounts to changing the input. Bringing down the input contrast is equivalent to shifting the contrasts-response function rightwards. We hypothesized that the suppression of optokinesis could be achieved mostly through response gain modulation, with little attenuation of the sensory input (rightwards shift).

On the other hand, if suppression occurs at the level of the input (contrast gain modulation), we should see a rightward shift in the contrast-response function (Figure 1B), as higher contrasts are needed to elicit the same response. This could be the case if, for instance, a signal originating in pre-motor areas selectively inhibits neurons that process motion in the reafferent direction and drive pursuit, such as those found in area MST, known to combine retinal and extra-retinal pursuit signals^21^. There are also reports consistent with sensory suppression for motion in the reafferent direction during pursuit, such as reports regarding lower contrast sensitivity^22^ or suppressed motion blur—accounted by a sharpening of the temporal impulse response function—in the reafferent direction^23,24^.

To tell whether optokinetic response during pursuit fit a response gain, sensory attenuation (a.k.a. contrast gain) or both accounts, we measured ocular tracking response to a background motion perturbation of varying contrasts. Strikingly, we found that the Naka-Rushton function does not provide a full description of oculomotor responses to motion in the reafferent direction. The most parsimonious model requires an extra parameter, corresponding to a contrast-dependent gating mechanism that would selectively cancel responses to high contrast signals.

## Methods

### Participants

Participants (N=12, 19-42 years old, 7 females) were undergraduate and postgraduate students from the University of Leicester’s School of Psychology, in addition to three authors. They were naïve to the purpose of the experiment and received vouchers for their participation of a value of £10 per hour. The experimental protocol was approved by the University of Leicester School of Psychology and Vision Sciences Ethics Committee, conforming with the guidelines of the Declaration of Helsinki. The study was not pre-registered.

### Materials

A CRT screen was used (Hewlett-Packard P1130, 1280 x 1024 pixels, 85 Hz) to display stimuli 60 cm from the participant. We used a chinrest to keep the head in place. The right eye (pupil centroid) was tracked by using a video-based eye-tracker (Eyelink 1000, SR Research Ltd, Osgoode, Ontario, Canada) at 1000 Hz. The stimuli were generated by using the Psychophysics toolbox PB-3 for MATLAB^25,26^. The stimuli were gamma corrected and displayed in gray scale at a 16-bit depth by using a DATAPixx Lite video processor (VPixx Technologies Inc., Saint-Bruno, Canada).

### Paradigm and procedure

To probe suppression of optokinesis we measured ocular responses in the direction of a brief and fast motion in the background during a pursuit eye movement^11,12,18–20^. We selected stimulation parameters that maximize responses to the background^18,27^. As shown in Figure 2A, the background consisted of low spatial frequency gratings (0.22 c/deg) oriented either vertically or horizontally, which covered the entire surface of the screen (32.08 deg x 25.82 deg), except for a narrow (1.03 deg) horizontal band (mid-gray, with a luminance of 46.19 cd/m^2^) in the center of the screen, over which the pursuit target moved horizontally. The background grating had one of seven contrast levels (1%, 2%, 4%, 8%, 16%, 32%, or 64%) and its phase was randomized on every trial between −π to π radians. Its maximal and minimal luminance was 0.03 and 92.01 cd/m2. A fixation dot (0.5 deg) stayed on the screen for 0.5 seconds at 5 deg either left or right off the screen center. Then the dot moved horizontally to the opposite side at a constant speed (10 deg/s) for 1 second. The participant’s task was to fixate and pursue the dot as accurately as possible, while ignoring events occurring in the background. The target direction (leftward if starting from the lefthand side of the screen, rightward otherwise) was randomized within blocks. In some trials the background drifted for a brief time (94 ms) at a high speed (57 deg/s) in the middle of the target trajectory, as shown in Figure 2B. The motion of the background could be upwards or downwards with the horizontal grating, and in the direction of pursuit (“With pursuit” condition) or opposite to the direction of pursuit (“Against pursuit” condition) with the vertical grating. There were catch trials for every background orientation and contrast, during which there was no motion of the background.

**Figure 2.**
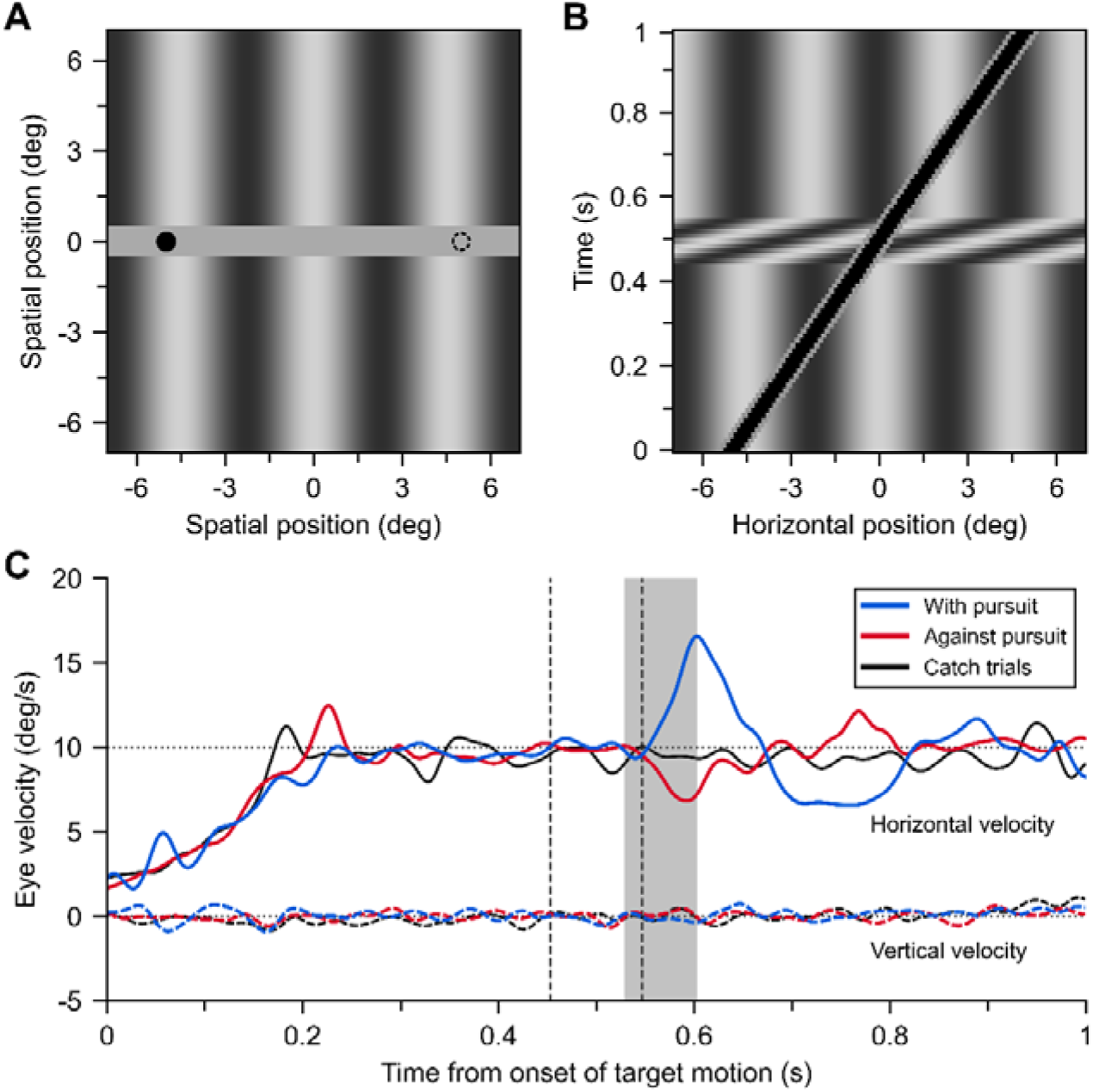
Example of the stimulus and data from the experiment. A) Example stimulus, where the target (black dot) appears on the left side of the screen. It moves for 1 second at 10 deg/s to the location indicated by the dashed circle. B) Space-time plot of the stimulus in a trial where the target moves to the right side of the screen and the background motion occurs in the direction of pursuit. C) Mean horizontal and vertical eye velocities of participant S11 to perturbations with and against pursuit, plus their corresponding catch trials. The contrast of the background grating was 4% (note that this contrast is lower than the ones shown in panels A and B). The vertical dashed lines indicate the temporal interval where the rapid background perturbation was introduced, and the shaded area corresponds to the interval that we considered for the analysis of the ocular tracking responses.1

We had then 7 contrasts by 6 conditions (“Upwards”, “Downwards”, “Against pursuit”, “With Pursuit”, “Catch horizontal”, “Catch vertical”). We had 5 sessions (252 trials), each with 6 repetitions per condition, leading to a total of 30 repetitions per condition across sessions (1260 trials).

Movies S1-4 (https://figshare.com/s/71391fc3a57e14e7cee3) illustrate the stimulation in four conditions (“Upwards”, “Downwards”, “Against pursuit”, “With Pursuit”).

### Data Analysis

Saccades and blinks were detected by using the Eyelink parser, using a 22 deg/s velocity and a 5000 deg/s^2^ acceleration threshold. This threshold was applied above a velocity baseline measured over the preceding 40 ms, up to 60 deg/s, to account for the ongoing pursuit eye movement. For further processing, we derived eye movement velocity by two-point differentiation of the eye position record, using a 20-ms step^28^. This signal was further filtered by applying a second-order low-pass Butterworth filter with a 20-Hz cutoff.

Figure 2C shows the mean horizontal and vertical eye velocities of one participant in response to the background motion in conditions “With pursuit”, “Against pursuit”, and catch trials. We plot here the average of de-saccaded and linearly interpolated velocity traces. For this purpose, we extended the Eyelink definition of saccade onsets and offsets by 30 ms based on visual inspection of velocity profiles. Our main analysis concerns a short period of time after background motion. This window is deliberately narrow^27^, within 75 to 150 ms post-motion onset, such that we can average pursuit eye movements uncontaminated by catch up saccades. We then average data in trials when no saccade was executed during this time-period. We obtained 54-100% (M=87%, SD=13%) of valid trials across participants based on the above criteria. To estimate the strength of the ocular tracking response we subtracted the catch trials’ velocity in the corresponding condition (a combination of contrast and background grating orientation) to the average velocity in the presence of background motion. We express responses in a reference frame that is relative to the direction of the background motion. Positives responses indicate an acceleration in the direction of the background motion, and negative responses a deceleration, relative to catch trials.

To fit the data, we used the Naka-Rushton function, as plotted in Figure 1A-B:

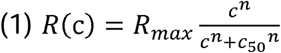

In Equation (1), *R_max_* is the maximum response without suppression, *c*_50_ is the contrast at half the maximum response, and *n* controls the steepness of the transition between minimum to maximum responses.

### Code accessibility

Code used for stimulus display, data analysis, and modelling will be made publicly available after publication (https://osf.io/5jm2r/).

## Results

We probed the suppression of optokinesis during pursuit eye movements by analysing reflexive responses to a brief background motion perturbation. Figure 2C shows average (de-saccaded and interpolated) velocity traces in one exemplary participant in different conditions. In this example, the pursuit target moves rightward, and a vertical background grating moves briefly (temporal interval indicated by dashed lines) either in the target direction (“With pursuit”, blue line), the opposite direction (“Against pursuit”, red line) or it does not move (“Catch trial”, black line). The background grating contrast is 4% in these examples. We can see that, relative to the catch trials eye movement velocity, after introducing the background perturbation, there is a deceleration of horizontal pursuit eye movements (reaching a velocity of about 7 deg/s) in the “Against pursuit” condition, but a much larger acceleration in the “With pursuit” condition (reaching a velocity of about 15 deg/s).

To better quantify the magnitude of the response in each condition (a combination of background motion direction and contrast), we calculated the average eye velocity over a 75-150-ms interval after the background perturbation is applied. Then, we subtracted their respective catch trials average (the same combination of background contrast and orientation) and expressed the response direction relative to background motion, with positive values representing a change in velocity in the direction of the background motion.

Figure 3 shows the group averages (mean ± s.e.m.) of the mean response to each perturbation direction as a function of the background’s contrast level. Figure 3A contains the horizontal eye responses to horizontal perturbations. When the perturbation occurred in the direction of pursuit, the horizontal eye response increased with increasing contrast levels, and then saturated at about 16% contrast. Perturbations opposite to the pursuit direction (“Against pursuit”), however, produced a very different response pattern. As expected, we have suppression, but we also see that this suppression, in addition to being consistent with response gain modulation (a change in Rmax), shows an unexpected increase with higher contrasts. Instead of the expected saturation, average responses decreased at 16% contrast to be virtually null at the highest contrasts. These results suggest that response in the reafferent direction does not follow the expected pattern and additional mechanisms, other than contrast or response gain modulation, as modelled by the Naka-Rushton function (black line), are necessary to fit the data.

**Figure 3.**
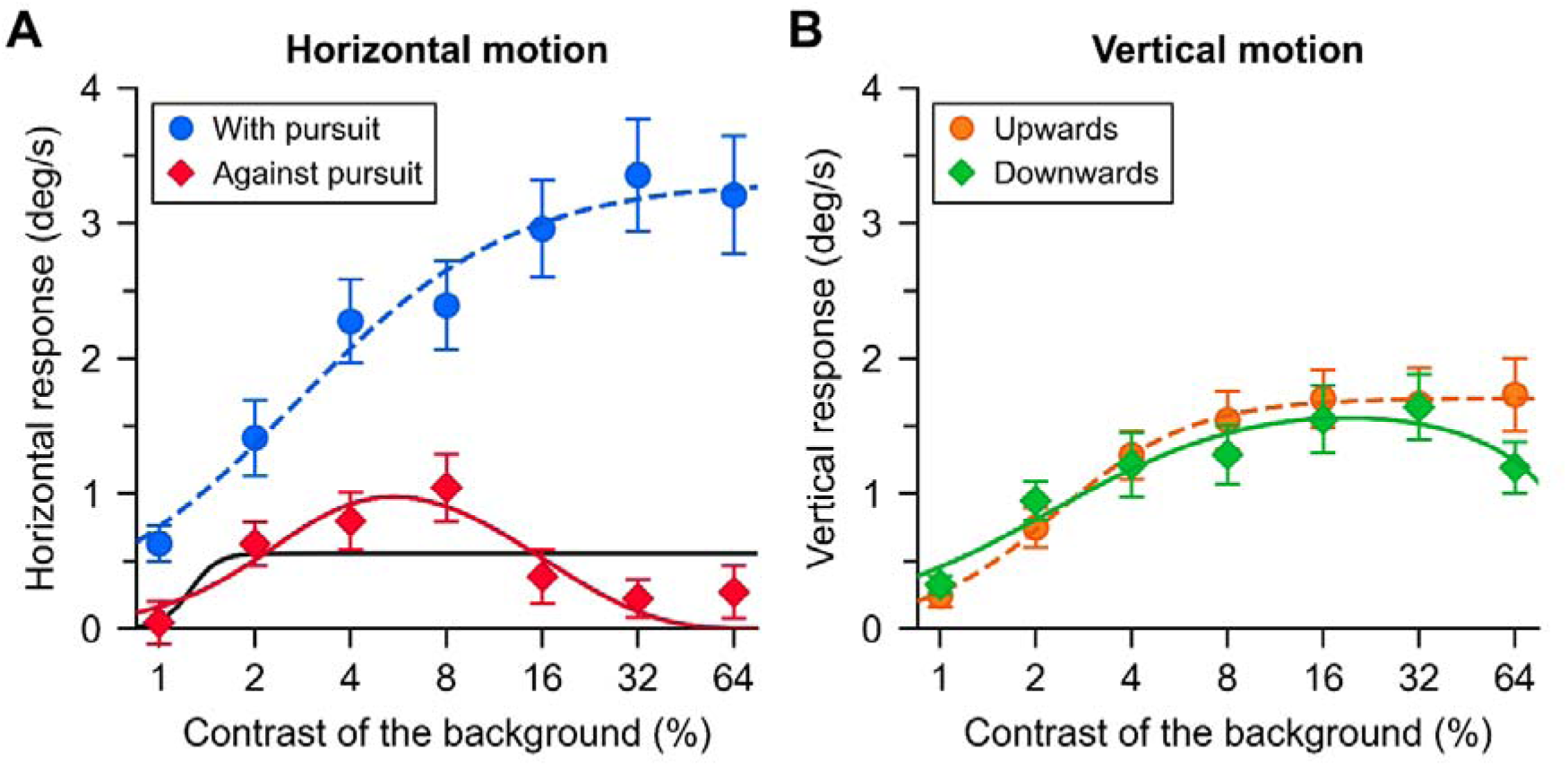
Group average (N=12) eye movement responses in the direction of background motion for the four perturbation directions as a function of the background contrast. A) Responses in the direction of pursuit (blue) and responses opposite to the pursuit direction (red). The solid black line shows the Naka-Rushton fit, whereas the solid red line and dashed blue line shows the fits of the gating model. B) Group average vertical eye movement responses to vertical motion and the fits of the gating model.

Figure 3B shows the vertical eye response to the vertical background perturbations. There were small differences between upwards and downwards perturbations at all contrast levels. Responses, as expected, increased with contrast, saturating at about 8% contrast.

There is a striking difference between these vertical responses, that is, responses orthogonal to the pursuit direction, and horizontal responses in the direction of pursuit. Besides seeing that both responses conform to the Naka-Rushton model, we observe that responses in the direction of pursuit reach much higher maximal values (“With pursuit” *R*_max_: 1.42-5.82 deg/s, average *R*_max_ of vertical directions: 0.69-2.77 deg/s), with a similar range of half-saturation values (“With pursuit” *c*_50_: 0.01-0.05 deg/s, average *c*_50_ of vertical directions: 0.01-0.03 deg/s). This change can be explained mostly by response gain modulation as suggested earlier^17^.

To assess whether response gain, sensory attenuation, or both explain the suppression of optokinesis in the reafferent direction, we fit a new model to the data that can accommodate the suppression observed in the “Against pursuit” condition. The black lines in Figure 3A show the fit of the Naka-Rushton function. Seeing that this function couldn’t account for the responses to perturbations opposite to the pursuit (Figure 3A), we fit a modified model in which there is an additional gating mechanism pinning down responses at higher contrasts:

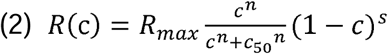

Equation (2) involves only the addition of one extra parameter to the Naka-Rushton function, *s*, controlling the intensity of the suppression at high contrasts. Because we observe complete suppression at the highest contrasts in the “Against pursuit” condition, we refer to it as a *gating* mechanism. Our intention at this stage is to provide a functional description. However, this description may encapsulate distinct mechanisms by which responses to reafferent motion are suppressed. The additional parameter, for instance, may represent the gating of a visuomotor response via stronger inhibitory connections between neural units that drive pursuit and neural units that are tuned to optic flow in the reafferent direction and high contrasts. The fits of the *gating model* (accounting for high-contrast gating in addition to response gain modulation, contrast gain modulation) appear in color in Figure 3A and 3B for each perturbation condition. We can see that this model accounts well for high-contrast suppression obtained in the condition “Against pursuit” (Figure 3A, red line).

The new model is better at the individual level as well. All participants show a similar pattern of responses in the reafferent direction (“With pursuit”), which we confirmed by comparing BICs based on squared residuals. Figure 4 shows the fit of the Naka-Rushton model and the gating model to the data of each participant. Starting with the condition “With pursuit”, the Naka-Rushton function appears as the best model for all but one participant. This is because, as we can see in Figure 4 (dashed lines and solid lines representing Naka-Rushton and gating-model fits, respectively), the fits for this condition are virtually the same in both models, but the BIC penalizes the gating model as it has one more parameter. In that condition, the suppression parameter is very close to 0 in all participants. On the other hand, in the condition “Against pursuit”, the gating model is superior for most participants. A close inspection of the fits in Figure 4 reveals that the suppression parameter is needed to better account for the suppression at higher contrasts.

**Figure 4.**
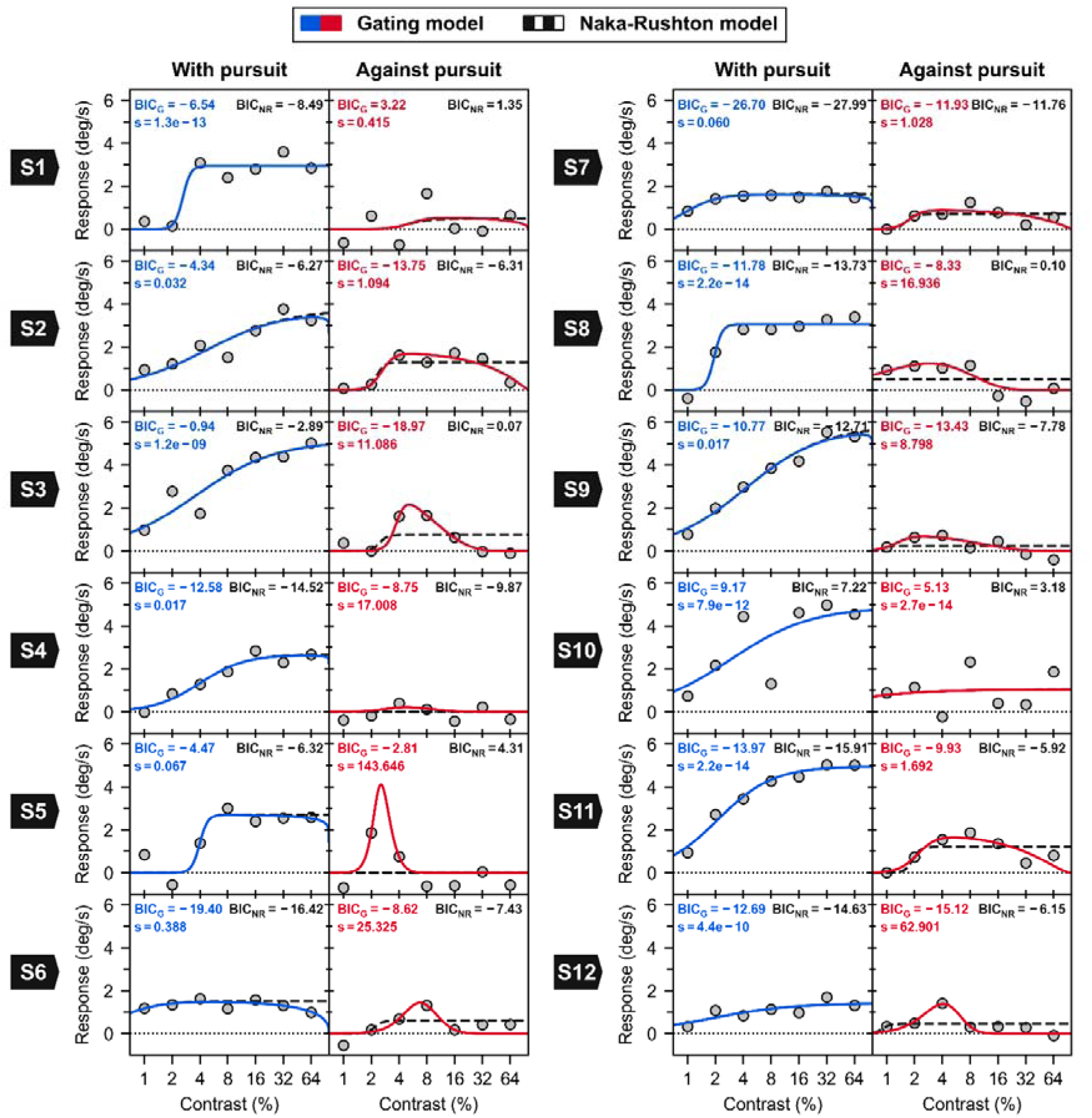
Fits of the gating and Naka-Rushton models to the data of the 12 participants for conditions with horizontal perturbation (i.e. With and Against pursuit). Colored lines correspond to the gating model, while dashed black lines correspond to the Naka-Rushton model.

After fitting the gating model to the data of each condition tested, we can now assess whether sensory attenuation (contrast gain modulation) does contribute to the suppression of optokinesis in addition to response gain modulation. If the fits show a decrease of *R*_max_ between conditions, while *c*_50_ remains constant, then this would suggest that a modulation of visuomotor gain could be responsible for the effect observed. On the contrary, an increase of *c*_50_, while *R*_max_ remains stable, would suggest that the suppression of optokinesis occurs because of sensory attenuation. Because there is a necessary extra parameter, *s,* in our model, we cannot directly compare *c*_50_, or *R*_max_ between “With pursuit” and “Against pursuit” to decide about which parameters are responsible for the suppression, since those co-vary with *s*. Instead, we compared the BICs for two models. A six-parameter model in which only *R_max_* and *s* are allowed to vary between conditions, but there is only one half-saturation (*c_50_*) parameter and *n* (slope) parameter—meaning that we fit different *R*_max_ and *s* for “Against pursuit” and “With pursuit” conditions, with a common *c*_50_ and n— and a seven-parameter model where *c*_50_ is also allowed to vary between conditions—meaning that we fit 3 parameters for “Against pursuit” (*R_max_*, *s*, *c*_50_) and 3 parameters for “With pursuit” (*R_max_*, *s*, *c*_50_), with a common *n*. The difference between the BIC values of both models is shown in Figure 5, while the fits and BIC values of each model appear in Table S1 and S2. The BIC values indicate that, with two exceptions where no model offers a very good fit to the data, a model where only response gain and *s* change between conditions provides a better and more parsimonious account of the data in most cases. Therefore, at the individual-level, there is little evidence that sensory attenuation is necessary to explain the data.

**Figure 5.**
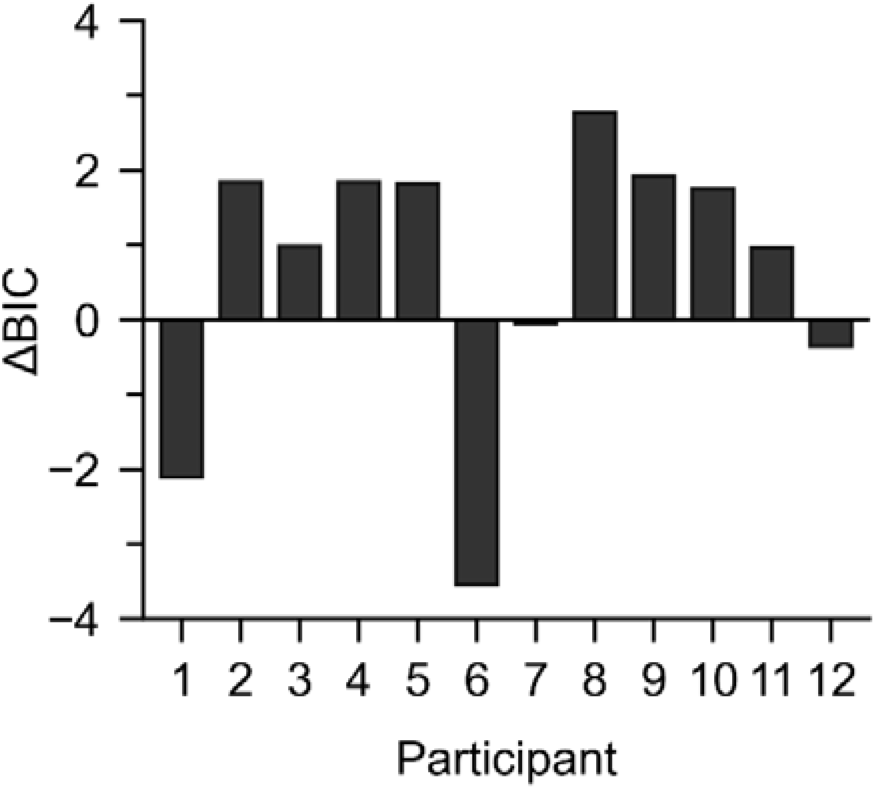
Difference in goodness of fit according to BIC criterion for the 7-parameter model, incorporating differences in half-contrast saturation depending on the direction of motion relative to pursuit-compared to the 6-parameter model for each participant. A positive difference indicates that the 6-parameter model is the better one

## Discussion

Our study aimed to differentiate between visuomotor response gain and sensory gain accounts of optokinesis suppression during smooth pursuit eye movements. Our findings suggest that neither account fully explains the observed reflexive eye movements in the reafferent direction. To adequately fit these responses, we introduced an additional parameter, that may correspond to an additional suppression: a high-contrast gating mechanism which could ensure undisturbed pursuit of a signal in the presence of highly salient background.

Key observations in our study are that:

1. Ocular tracking response in the perturbation direction were much larger in the pursuit direction compared to the vertical direction. This modulation maps well in a simple response gain modulation, as suggested earlier^29^.
2. Our results suggest that a third, newly observed, mechanism is necessary to explain suppression of optokinesis as probed by background motion, amounting to a cancellation of responses towards high contrast background motion opposite to the pursuit direction.
3. Our results suggest a limited role for sensory suppression. Instead, the suppression of response to background motion appears to be explained by a combination of visuomotor gain downregulation and high-contrast gating, or cancellation, since suppression is complete at the highest contrasts.

Several lines of evidence pointed to a visuomotor gain modulation mechanisms to regulate how pursuit eye movements adjust to visual feedback. A higher visuomotor gain during pursuit compared to fixation is necessary to ensure stable tracking in the presence of small retinal error signals. Single cell recordings provide evidence that the visuomotor gain, as is the visuomotor transformation in saccades, can be modulated in the frontal eye fields (FEF)^30^. FEF could provide a top-down signal by which the target is selected for pursuit eye movements and by which visuomotor gain is increased for signals in the direction of pursuit. Adaptation to stimulation statistics has been suggested to explain a wide range of adaptive behaviors^3,31^. This type of adaptive suppression requires a mechanism to feed information about movement as provided by a corollary discharge.

The frontal-eye-fields (FEF) have been proposed as a key neural substrate in the modulation of the internal visuomotor gain during pursuit^30^. This could explain several observed phenomena, including larger responses to target and background perturbations relative to fixation^29^, larger responses in the pursuit direction relative to the vertical direction^29^, and the suppression of optokinesis^30^. At the neural population level, we hypothesize that the high-contrast gating and visuomotor gain modulation could originate in FEF.

The exact level at which a high-contrast gating mechanism may operate is unclear. Supersaturation, observed in some neuronal responses, where maximal contrasts do not generate maximal responses, may play a role^32^. Contrast non-linearities can have a function in detecting visual features such as conjunctions. These non-linearities are much more common at the neuronal level in the cat and macaque cortex than previously thought. Neural contrast responses are also believed to be an adaptation to natural image statistics^33^.

In the presence of supersaturating responses to contrast, we can imagine high-contrast gating as arising from adaptative suppression to the contrasts that are prevalent in the background of natural images. An agent^34^ that would be rewarded by a sharper image when tracking a small target over a cluttered background, would need to suppress eye movements in the direction of the reafferent motion. If these responses are driven by motion signals carried by neurons with supersaturating responses to contrast, the backpropagation of error would suppress response to the prevalent scene contrasts but spare response to units that have maximal responses at lower contrasts. Natural image statistics suggest that while very low contrasts can occur over small patches, they are much less represented over large ones^35^. The global optic flow generated by pursuit eye movements, which processing could be suppressed in the reafferent direction, can be represented by neuronal units with large receptive fields, such as those found in the MST^36^.

Whereas Linder and Ilg refer to the suppression of optokinesis when measuring reflexive responses to the background, others have measured the optokinetic response (oscillating quick and slow eye movement beat) elicited after a period of stimulation^37–40^. Further studies could test whether these two paradigms show a consistent set of mechanisms.

Sensory attenuation of reafferent signals is observed across sensory modalities, for instance, in auditory neuron’s response to self-generated song in birds^3^ and other species. The attenuation of expected signals can participate to efficient coding of sensory information, whereby computing resources are preferentially allocated to novelty^41–44^. Expectations concern stimulus characteristics in interaction with the characteristics of goal-oriented behavior, including how visual stimulation is shaped by the constant eye movements we make^9,45^. However, we find little role of sensory attenuation, as suppression of the response to background motion is explained by downregulation of the visuomotor gain in the form of a response gain modulation combination with high-contrast gating. One explanation is that although we observe sensory attenuation in motion integration tasks, which would suggest sparing of computation in the reafferent direction^10^, the processing of reafferent signals may also have an important role in recalibrating motion perception in spatial coordinates^8^.

We show that the suppression of reflexive oculomotor responses to background motion (optokinesis) in the reafferent direction during pursuit eye movements does not conform to a simple response gain modulation, contrast gain modulation, or any combination of contrast and response gain modulation, but that it includes a high-contrast gating mechanism, which could have arisen through adaptation to natural image statistics.

## Acknowledgements

Supported by grant PDI2021-122245NB-I00 from Ministerio de Ciencia e Innovación (Spain) to ISP. For the purpose of open access, the author has applied a Creative Commons Attribution license (CC BY) to any Author Accepted Manuscript version arising from this submission.

---

**Table S1.** Best fits to individual data based on a 6-parameter functional model. In this model, the maximal response, *R*_max,_ and suppression parameter, *s,* depend on the direction of background motion relative to the pursuit eye movement’s direction (4 parameters), while *n* and the half-saturation contrast parameter, *C*_50_, are direction independent. The root of the sum of the squared residuals (RSS) and the Bayesian Information Criterion (BIC) are shown for this model.

| | $R_{\max}$. With | $R_{\max}$. Against | $n$ | $C_{50}$ | $s_{With}$ | $s_{Against}$ | RSS | BIC |
| --- | --- | --- | --- | --- | --- | --- | --- | --- |
| S1 | 2.94 | 0.32 | 21.73 | 0.02 | 0.00 | 0.00 | 4.98 | 9.30 |
| S2 | 13.17 | 9.78 | 0.62 | 1.00 | 2.15 | 0.53 | 1.65 | 1.55 |
| S3 | 5.20 | 2.91 | 1.01 | 0.04 | 6.14 | 0.00 | 3.37 | 6.56 |
| S4 | 2.64 | 0.47 | 1.85 | 0.04 | 19.27 | 0.00 | 1.10 | -1.26 |
| S5 | 2.70 | >100 | 12.07 | 0.04 | 410.94 | 0.06 | 2.79 | 5.23 |
| S6 | 1.61 | 0.50 | 2.16 | 0.01 | 0.14 | 0.49 | 1.80 | 2.15 |
| S7 | 1.66 | 0.83 | 2.79 | 0.01 | 0.73 | 0.09 | 0.64 | -5.11 |
| S9 | 3.13 | 3.13 | 3.09 | 0.02 | 17.87 | 0.00 | 1.93 | 2.66 |
| S10 | 5.72 | 2.09 | 1.07 | 0.04 | 13.49 | 0.01 | 0.91 | -2.64 |
| S11 | 14.01 | 3.31 | 0.45 | 0.98 | 0.00 | 0.30 | 13.46 | 16.25 |
| S12 | 4.88 | 1.99 | 1.71 | 0.02 | 1.84 | 0.00 | 1.20 | -0.67 |

**Table S2.**
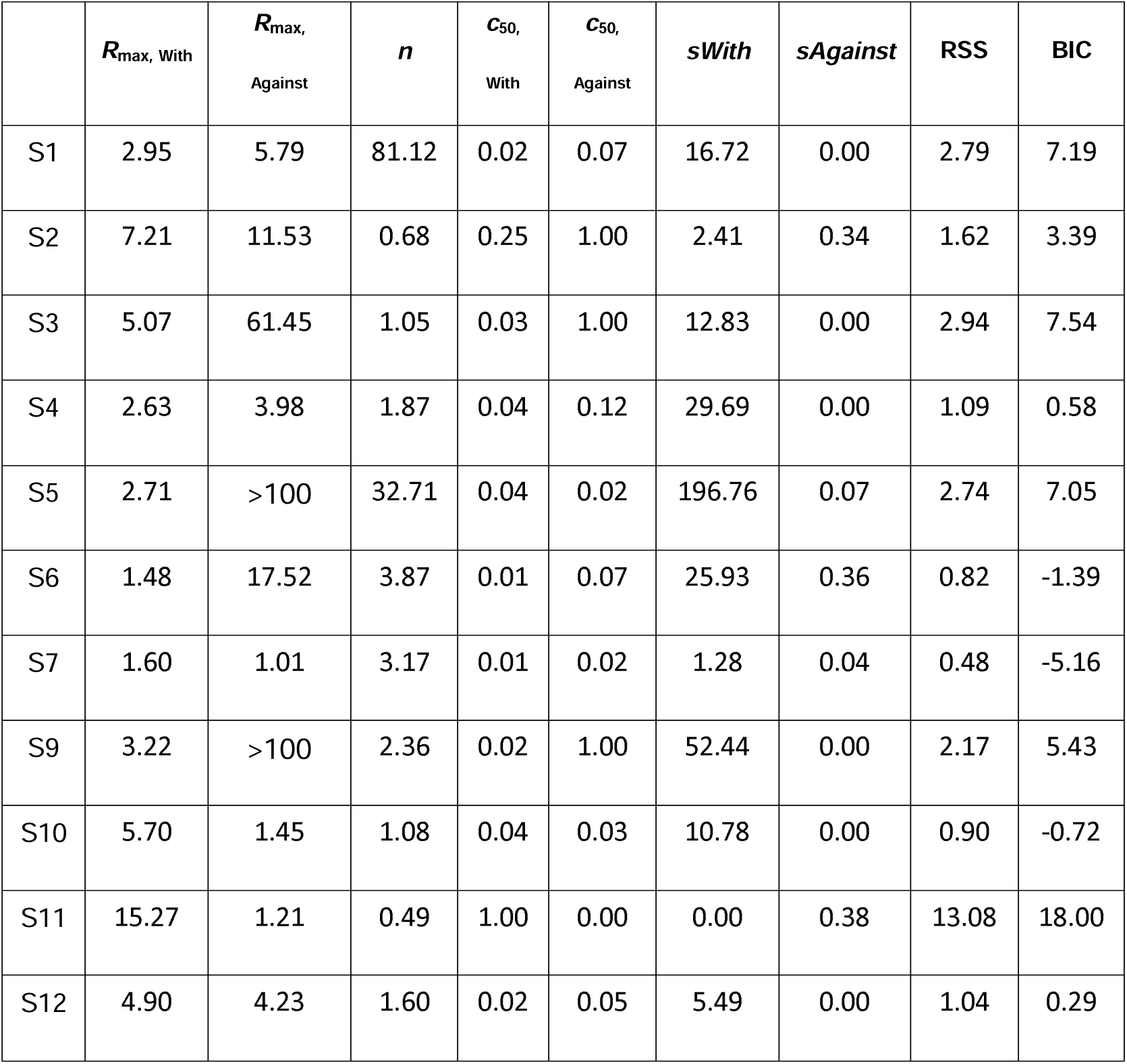
Best fits to individual data based on 7-parameter functional model. In this model, the maximal response, *R*_max_, suppression parameter, *s,* and the half-saturation contrast parameter, *C*_50,_ depend on the direction of background motion relative to the pursuit eye movement’s direction (6 parameters), and *n*. The root of the sum of the squared residuals (RSS) and the Bayesian Information Criterion (BIC) are shown for this model.

| | $R_{\max}$, With | $R_{\max}$,<br>Against | $n$ | $C_{50}$,<br>With | $C_{50}$,<br>Against | $s$With | $s$Against | RSS | BIC |
| --- | --- | --- | --- | --- | --- | --- | --- | --- | --- |
| S1 | 2.95 | 5.79 | 81.12 | 0.02 | 0.07 | 16.72 | 0.00 | 2.79 | 7.19 |
| S2 | 7.21 | 11.53 | 0.68 | 0.25 | 1.00 | 2.41 | 0.34 | 1.62 | 3.39 |
| S3 | 5.07 | 61.45 | 1.05 | 0.03 | 1.00 | 12.83 | 0.00 | 2.94 | 7.54 |
| S4 | 2.63 | 3.98 | 1.87 | 0.04 | 0.12 | 29.69 | 0.00 | 1.09 | 0.58 |
| S5 | 2.71 | >100 | 32.71 | 0.04 | 0.02 | 196.76 | 0.07 | 2.74 | 7.05 |
| S6 | 1.48 | 17.52 | 3.87 | 0.01 | 0.07 | 25.93 | 0.36 | 0.82 | -1.39 |
| S7 | 1.60 | 1.01 | 3.17 | 0.01 | 0.02 | 1.28 | 0.04 | 0.48 | -5.16 |
| S9 | 3.22 | >100 | 2.36 | 0.02 | 1.00 | 52.44 | 0.00 | 2.17 | 5.43 |
| S10 | 5.70 | 1.45 | 1.08 | 0.04 | 0.03 | 10.78 | 0.00 | 0.90 | -0.72 |
| S11 | 15.27 | 1.21 | 0.49 | 1.00 | 0.00 | 0.00 | 0.38 | 13.08 | 18.00 |
| S12 | 4.90 | 4.23 | 1.60 | 0.02 | 0.05 | 5.49 | 0.00 | 1.04 | 0.29 |

## Notes

https://osf.io/5jm2r/

https://figshare.com/s/71391fc3a57e14e7cee3

### Competing Interest Statement

The authors have declared no competing interest.

